# Blood meals from ‘dead-end’ vertebrate hosts enhance transmission potential of malaria-infected mosquitoes

**DOI:** 10.1101/2022.12.24.521872

**Authors:** Ashutosh K. Pathak, Justine C. Shiau, Cury Rafael Sadock de Freitas, Dennis E. Kyle

## Abstract

Ingestion of an additional blood meal(s) by mosquitoes can accelerate parasite migration to the salivary glands in infected mosquitoes. Most studies, however, offer blood from the same vertebrate host species as the original challenge (for e.g., human for primary and additional blood meals). Here, we show a second blood meal from bovine and canine hosts can also enhance sporozoite migration in *Anopheles stephensi* mosquitoes infected with the human- and rodent-restricted *Plasmodium falciparum* and *P. berghei* respectively. The extrinsic incubation period (time to sporozoite appearance in salivary glands) showed more consistent reductions with blood from human and bovine donors than canine blood, although the latter’s effect may be confounded by the toxicity, albeit non-specific, associated with the anticoagulant used to collect whole blood from donors. The complex patterns of enhancement highlight the limitations of a laboratory system but are nonetheless reminiscent of parasite host-specificity and mosquito adaptations, and the genetic predisposition of *An. stephensi* for bovine blood. We suggest that in natural settings, a blood meal from any vertebrate host could accentuate the risk of human infections by *P. falciparum*, although our observations should also be applicable to other species of *Plasmodium* and indeed other vector-borne diseases such as arboviruses for instance.

**Graphical Abstract:** 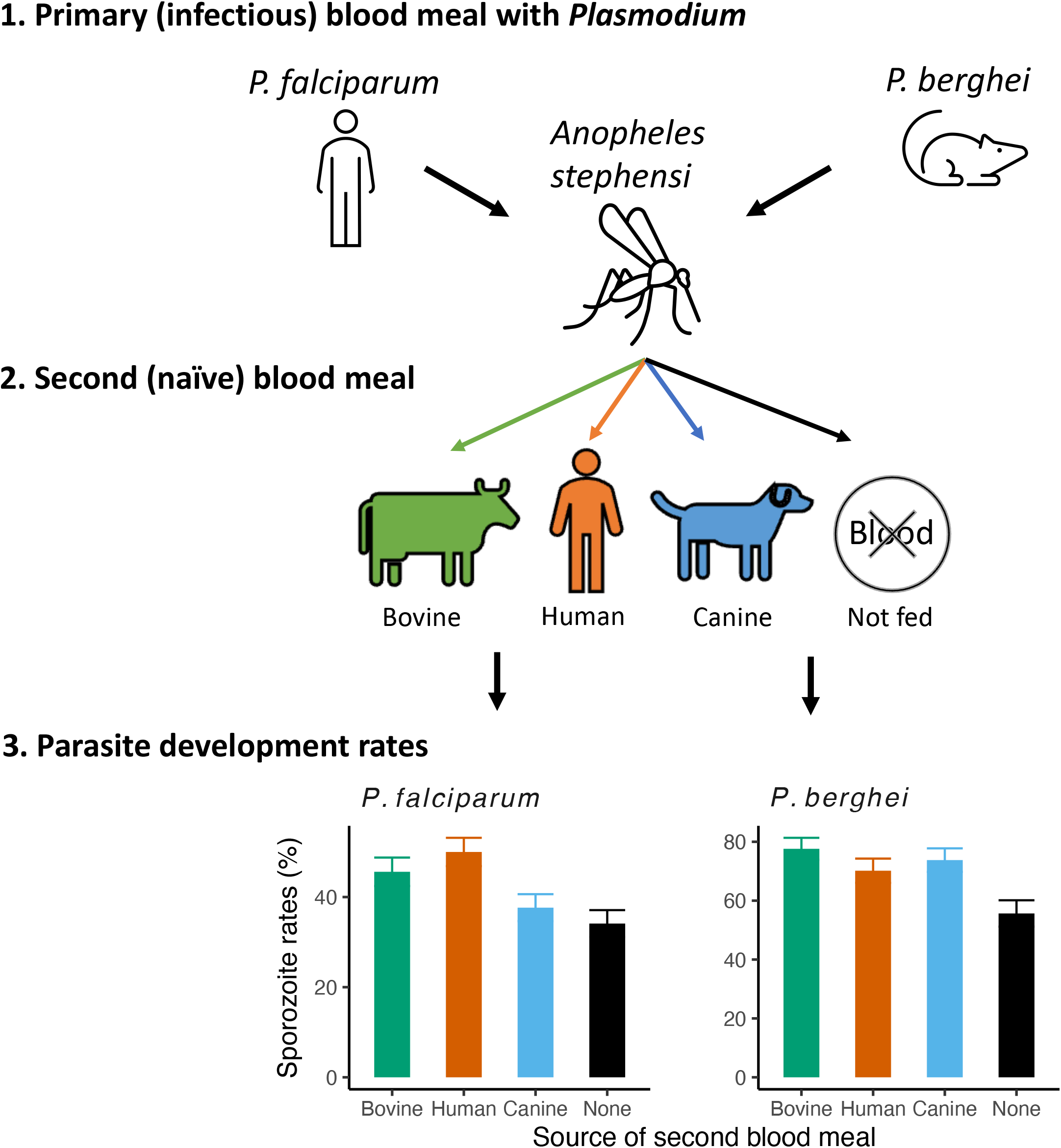

**HIGHLIGHTS:**

1. *Plasmodium falciparum* and *P. berghei* cause malaria in humans and rodents, respectively.
2. They are vectored by *Anopheles* species that can also imbibe blood from other vertebrate species.
3. Parasite development is enhanced after another blood meal, usually from the same host (e.g., human).
4. Here, bovine, and canine blood also enhanced development of *Plasmodium* species in *An. stephensi*.
5. In natural settings, blood from any host could enhance development of vector transmitted parasites.

## INTRODUCTION

Hematophagy is central to the survival of mosquitoes, but also the transmission of several diseases. While parasite transmission requires two independent biting events, in general, mosquitoes can acquire multiple blood meals over their lifetime [1]. Ingestion of an additional blood meal(s) has been shown to enhance vectorial capacity of *Plasmodium*- and arbovirus-carrying *Anopheles* and *Aedes* species of mosquitoes respectively, by accelerating rates of parasite replication and migration to the salivary glands (i.e., shortening the extrinsic incubation period (EIP)) in addition to promoting vector survival [2-5]. However, in most of these studies, mosquitoes were offered additional blood meals from the same host species as the original challenge (e.g., human for primary and additional blood meals) (reviewed in [3] but see [6]). While proximity to livestock is a well-known risk factor for malaria incidence, surveillance of natural populations of mosquitoes routinely finds evidence of blood from multiple host species while co-incident with human *Plasmodium* infections [7-13]. However, it is generally assumed that the nutritional benefits of a non-human blood meal can enhance vectorial capacity by promoting vector survival and population density [14-20], although it is unclear if it can influence parasite development rates directly. If non-human blood meals were to enhance parasite development rates, we may need to consider non-human hosts more closely as an additional risk factor directly impacting parasite transmission.

The importance of non-human blood meals is highlighted by the declining effectiveness of insecticide-treated bednets (ITNs) in sub-Saharan Africa [21]. The reduced effectiveness is associated with the emergence of insecticide-resistant *Anopheles*, as well as selection for altered feeding behavior amongst mosquito species [15,17,22]. Altered feeding behavior can range from shifts in time when mosquitoes bite to the recognition of mosquitoes with mixed host preference that can still vector human parasites. This mixed host preference constitutes the rationale for administering endectocides to companion animals for malaria control, and the One Health initiative in general [18,20,23]. However, identifying potential sources of variation and the direct consequences for parasite transmission are also important.

In the current study, we provide rationale for two such sources of variation, and then proceed to test them empirically. First, mosquito species may differ in how well they can digest and assimilate nutrients from the blood of different vertebrate host species [1,7,24], and in turn, the amount of nutrition available to the parasite; in other words, blood meals from different host species are likely to benefit the parasite differently. Second, although the same *Plasmodium* species (e.g., *P. falciparum*) can be vectored by several *Anopheles*, they generally only infect a single vertebrate host species (humans); as such, it is possible this parasite benefits more from human blood than a non-human blood meal. In the current study, *An. stephensi* mosquitoes were infected with the human-specific *P. falciparum* or the rodent-restricted *P. berghei* and offered an additional blood meal from human, bovine, canine and murine donors. Changes in vectorial capacity were assessed by measuring the extrinsic incubation period (EIP, defined as the days post-blood meal or time when sporozoites are first observed in the salivary glands) as well as the probability of daily survival [2]. We asked, 1) how blood from phylogenetically distant (but ecologically related) hosts with different biochemical compositions alter vectorial capacity, and 2), whether this effect was dependent on *Plasmodium* host specificity. We find that in general, an additional blood meal enhanced vectorial capacity irrespective of the source suggesting that any non-human blood meal could enhance the vectorial capacity of mosquitoes infected with any species of *Plasmodium*. However, our results suggest caution in interpreting host specificity while also highlighting the limitations of extrapolating mosquito behavior from a laboratory system.

## MATERIALS AND METHODS

### Study design

On the day of infection (0 days post-infection), 1250 female *An. stephensi* were infected with *P. falciparum* NF54 or *P. berghei* (PbGFP-LUC_CON_) respectively, as described previously [25-27]. At 6 days post-infection, oviposition sites were introduced into the mosquito cages infected with either parasite. Mosquitoes were allowed to oviposit over two and six days for *P. falciparum-* and *P. berghei*-infected mosquitoes, respectively, based on the temperature-specific gonotrophic cycles for *An. stephensi*, as shown previously [28]. Prior to being separated into the respective cages in preparation for the second blood meal at 8 and 12-days post-infection for *P. falciparum* and *P. berghei* infected mosquitoes respectively, a subset of individuals in each replicate (n=40-41) were used to determine the gravid status (presence or absence of eggs in ovaries, also see below), infectivity of the initial infectious blood meal (presence or absence of oocysts in the midguts), and if any of these individuals’ salivary glands were already sporozoite-positive (Supplementary table 1). Next, ∼200 mosquitoes were sorted into each of four new cages. One cage was not offered a second blood meal (control, or ‘none’ herein) while the remaining three experimental cages were offered a second blood meal of bovine, canine or human origin respectively. After the blood meal, 20-25 mosquitoes from each group were sampled at regular intervals, with salivary glands and ovaries (see below) assessed for sporozoite and gravid status respectively, as described previously [25-27]; for *P. falciparum*, mosquitoes were sampled on 10, 12, 14, 16, 18, and 21 days post-infection (or 2, 4, 6, 8, 10 and 13 days post-second blood meal), while *P. berghei*-infected mosquitoes were assessed at 14, 16 and 18 days post-infection (or 2, 4 and 6 days post-second blood meal).

As mentioned above, in addition to checking salivary glands for the presence or absence of sporozoites in the salivary glands, ovaries of all mosquitoes were also assessed for gravidity (presence or absence of eggs). While the former measure was to estimate the EIP, the latter measure was to ensure our results fulfilled two criteria: 1) mosquitoes should have oviposited prior to the second blood meal (‘nulliparous’ or not gravid), such that 2), any changes in gravid status after this blood meal would be reflective of feeding rates in general, but also if the mosquitoes preferred blood from a particular vertebrate species over another. Since the control group (not offered a second blood meal) was representative of the starting mosquito populations for all the groups (Supplementary figure 1), rates of oviposition before the second blood meal were extrapolated from the gravid status of mosquitoes in the control groups; ideally, high rates of nulliparity/non-gravid status (absence of eggs in ovaries) would indicate that mosquitoes had successfully oviposited prior to being sorted for the second blood meals. By contrast, high rates of parity/gravid status (presence of eggs in ovaries) in the experimental groups would provide an overview of feeding rates with the second blood meal, with any differences in the proportions of gravid mosquitoes between the groups indicative of the mosquito colony’s preference for one source of blood over another: no clear differences in gravid status between the various species’ blood meals would confirm the lack of preference. Note that repeated measurements of gravid status from different individuals over time in the respective groups would also impart more robustness to the results [25].

In general, our results mostly describe the effect of additional blood meals over two independent replicates for mosquitoes infected with either parasite, with each replicate defined based on using two independent donors corresponding to the respective host species. Note that in the first two replicates with *P. berghei*-infected mosquitoes, we tested human, bovine, and canine blood from the same two donors used to assay *P. falciparum*-infected mosquitoes above (Figure 2); in the third replicate, the effect of mouse and human blood on sporozoite migration rates was assessed with an independent set of donors (Supplementary figure 1). Sporozoite invasion rates in the first two replicates were estimated at 14-, 16- and 18-days post-infection (or 2, 4 and 6 days after the second blood meal at 12 days post-infection) and 16-, 18- and 20-days post-infection in the third replicate.

### Parasites

*P. falciparum* NF54 was propagated in vitro as detailed previously [27]. *P. berghei* was propagated in mice as described previously [25].

**Table 1.** Statistical modeling of sporozoite rates for groups challenged with either parasite, to proportions of mosquitoes with sporozoites in the salivary glands from total sampled.

| *Predictors* | *P. falciparum* sporozoite rates |  |  | *P. berghei* sporozoite rates |  |  |
| --- | --- | --- | --- | --- | --- | --- |
|  | *Odds Ratios* | *CI* | *p* | *Odds Ratios* | *CI* | *p* |
| (Intercept) | 0.50 | 0.19 – 1.32 | 0.161 | 1.26 | 0.88 – 1.80 | 0.202 |
| Bovine blood [vs. None] | 2.80 | 1.52 – 5.17 | **0.001** | 2.95 | 1.67 – 5.21 | **<0.001** |
| Human blood [vs. None] | 4.56 | 2.50 – 8.32 | **<0.001** | 2.02 | 1.17 – 3.50 | **0.012** |
| Canine blood [vs. None] | 2.12 | 1.17 – 3.84 | **0.013** | 2.24 | 1.31 – 3.85 | **0.003** |
| Days post-infection (dpi) | 22.86 | 12.03 – 43.46 | **<0.001** | 1.15 | 0.81 – 1.64 | 0.433 |
| Dpi<sup>2</sup> (to model 'quadratic/humped' migration rates) | 0.25 | 0.19 – 0.34 | **<0.001** |  |  |  |
| Bovine blood * dpi | 0.83 | 0.41 – 1.66 | 0.594 | 1.38 | 0.78 – 2.45 | 0.266 |
| Human blood * dpi | 0.54 | 0.28 – 1.05 | 0.068 | 1.75 | 1.01 – 3.02 | **0.046** |
| Canine blood * dpi | 0.39 | 0.21 – 0.74 | **0.004** | 1.08 | 0.63 – 1.85 | 0.793 |
| Replicates | 2 |  |  | 2 |  |  |
| Observations | 1022 |  |  | 495 |  |  |

### Mosquitoes

Our *An. stephensi* colony originates from the Walter Reed Biosystematics Unit where it has been maintained since 1961. Since 2015 however, we have been maintained the colony solely on human blood. General husbandry was performed as detailed previously [25-27].

### Preparation of various blood meals

Human blood (packed RBCs) and plasma were purchased from a commercial supplier (Interstate Blood Bank, Asheville, NC). Bovine and canine blood was provided by Dr. Kelsey Hart and Dr. Andrew Moorhead of the University of Georgia Veterinary School respectively. Licensed technicians collected whole blood in heparin or Ethylenediaminetetraacetic acid (EDTA) to prevent coagulation. Animals were maintained in SPF conditions and had never been dosed with anthelminthics or endectocides.

For all blood meals, immediately after blood collection, red blood cells and plasma were separated by centrifugation at 1800 g for 5 minutes at room temperature. Packed red blood cells were stored at 4°C and plasma was heat-inactivated at 56°C for 30 minutes in a water bath and then stored at -20°C. Just prior to being offered to mosquitoes, packed RBCs from either source was reconstituted to ∼50% hematocrit with freshly thawed donor-matched plasma. Refrigerated, packed RBCs from all sources of blood were less than two weeks post-collection, as suggested previously [27].

### Mosquito infections and assessment

3 to 5-day old female *An. stephensi* were infected as described in detail previously for *P. falciparum* [26,27] and *P. berghei* [25]. To stimulate oviposition at 6 days post-infection, mosquitoes were offered open-top polypropylene cups (4.5 inches diameter x 2.5 inches height) containing ∼200ml deionized water. Parasite densities and infection status in the mosquitoes were estimated at the stated time points as described before [25-27]. Mortality was assessed daily in all the cages.

### Data analysis

Data analysis and modeling was performed in the RStudio IDE (version 2022.07.1, Build 554) running R software (version 4.2), as described in detail previously [25-27]. Unless stated otherwise, blood feeding rates and parasite prevalence (EIP) were modeled as the proportion gravid or infected from total dissected respectively, with a binomial distribution in generalized linear mixed effects models (GLMMs). Mosquito survival over time was modeled with mixed effects Cox proportional hazards in the ‘coxme’ package. Source species of blood meal and where applicable, donor or cage ID, were specified as categorical variables. Time was specified as a continuous variable after centering with the mean and scaling with standard deviation respectively, as described previously. Intercepts of the effects of blood species and/or time on parasite prevalence, feeding rates or survival were allowed to vary randomly between the replicates. Pairwise comparisons were performed post-hoc with the estimated marginal means from the various models, using Tukey’s method to adjust for multiple comparisons, as recommended by the ‘emmeans’ package.

## RESULTS

### Blood meals from distant vertebrate hosts and feeding rates of *Plasmodium*-infected mosquitoes

After the initial infectious blood meal, *P. falciparum*-exposed *An. stephensi* mosquitoes were provided oviposition sites at 6 days post-infection prior to the second blood meal. The proportion of mosquitoes still carrying eggs (gravid) in the control group (‘None’) did not change over the 10-21 days (Odd ratio (OR) = 1.13, 95% confidence intervals (95% CI) = 0.64 – 1.99, p=0.67) (Supplementary table 1), with an average of ∼5% of individuals in this group still parous (Supplementary figure 2a). Assuming a gravid mosquito after the second blood meal was indicative of a successful blood feed in the experimental groups, mean rates of gravidity in the latter groups (>75%) were clearly higher than the non-blood fed control group (∼5%) (Supplementary figure 2a). However, the proportion of gravid mosquitoes declined slightly over 10-21 days in the groups offered human blood meals (OR = 0.44, 95%CI = 0.23 – 0.84, p=0.013), and to a less clear extent canine blood (OR = 0.56, 95%CI = 0.29 – 1.04, p=0.068) (especially after 16 days post-infection), but not bovine blood (OR = 0.59, 95%CI = 0.29 – 1.04, p=0.11) (Supplementary figure 2a) (Supplementary table 1). Despite the declining rates of gravidity in the two former groups, pairwise comparisons of the means estimated from the model did not indicate any differences in the proportion of gravid (i.e., blood fed) mosquitoes between any group, at any sampling point (Supplementary table 2).

While *An. stephensi* infected with the rodent-specific *P. berghei* did not appear to show any changes in gravidity over time for any group (Supplementary figure 2b), gravidity was clearly reduced in the control (‘None’, mean = 11.3%) compared to the experimental groups (Supplementary table 1). Pairwise comparisons of the model-estimated means did not suggest any apparent differences in gravidity over time between mosquitoes offered bovine, canine, or human blood meals (Supplementary table 3). Taken together, these results suggest most *Plasmodium*-infected mosquitoes had successfully oviposited prior to the second blood meals; additionally, by showing how most mosquitoes in the groups offered second blood meals were able to make eggs irrespective of blood source (or donor), this result indicates a lack of preference for a particular host species (for rationale, see ‘Study design’ section under ‘Methods’, and Supplementary figure 1 for visual summary).

### Blood meals from distant vertebrate hosts and sporozoite development rates in *Plasmodium*-infected mosquitoes

In case of *P. falciparum* infected mosquitoes, except for canine blood, no group-specific differences were noted in the rates of migration, with the initial increase in sporozoite migration rates (OR = 22.86, 95%CI = 12.03 – 43.46, p<0.001) in all remaining groups eventually approaching saturation by 21 days post-infection (OR = 0.25, 95%CI = 0.19 – 0.34, p<0.001) (Figure 1a) (Table 1). In case of canine blood however, the initial enhancement in parasite development and migration was soon followed by a clear decline in the number of sporozoite-positive mosquitoes starting 16-18 days post-blood meal (OR = 0.39, 95%CI = 0.21 – 0.74, p=0.004) (blue lines, Figure 1a). This decline was likely why, compared to the groups that were not offered a second blood meal (“None”) canine blood showed the lowest enhancement in mean sporozoite rates (OR = 2.21, 95%CI = 1.17 – 3.84, p=0.013), with higher levels noted for bovine blood (OR = 3.06, 95%CI = 1.34 – 6.95, p=0.008) but especially in the groups offered human blood (OR = 5.43, 95%CI = 2.38 – 12.38, p<0.001) (Figure 1b).

**Figure 1.**
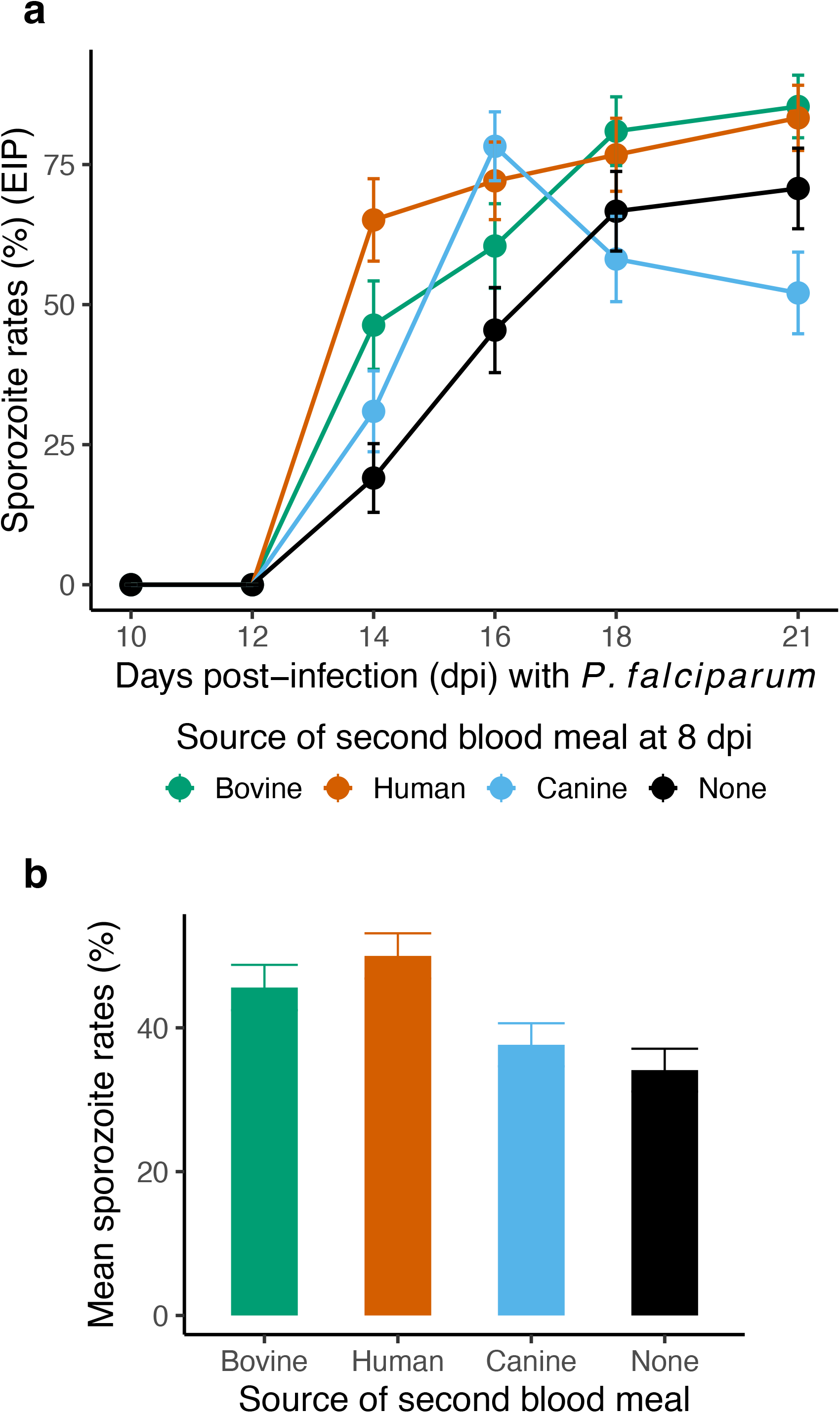
After the initial infectious blood meal, a second blood meal at 8 days post-infection from bovine (green lines/points/columns), human (red) and canine (blue) donors (**a**) enhances migration of human-specific *P. falciparum* sporozoites to the salivary glands (i.e., shorter extrinsic incubation period, EIP), and (**b**), the mean proportion of infected mosquitoes, compared to mosquitoes that were not offered a second blood meal (‘None’, black lines/columns). Data is presented as the mean and standard errors in the proportion of sporozoite-positive mosquitoes from total sampled over two independent infections (biological replicates); in each infection, whole blood from two independent donors from all sources was offered to the mosquitoes. For statistical analyses, refer to Table 1.

The effect of additional blood meals was also tested with mosquitoes infected with the rodent-restricted *P. berghei* (Figure 2) (Table 1). While sporozoite migration rates showed a marginal increase over time in the groups offered human blood (OR = 1.75, 95%CI = 1.01 – 3.02, p=0.046) (Figure 2a), in general, compared to the control groups (‘None’), the overall (mean) proportion of mosquitoes positive for *P. berghei* sporozoites were highest in the group offered bovine blood (OR = 2.95, 95%CI = 1.67 – 5.21, p<0.001), followed by canine blood (OR = 2.24, 95%CI = 1.31 – 3.85, p=0.003), and lastly, human blood (OR = 2.02, 95%CI = 1.17 – 3.5, p=0.012) (Figure 2b) (Table 1).

**Figure 2.**
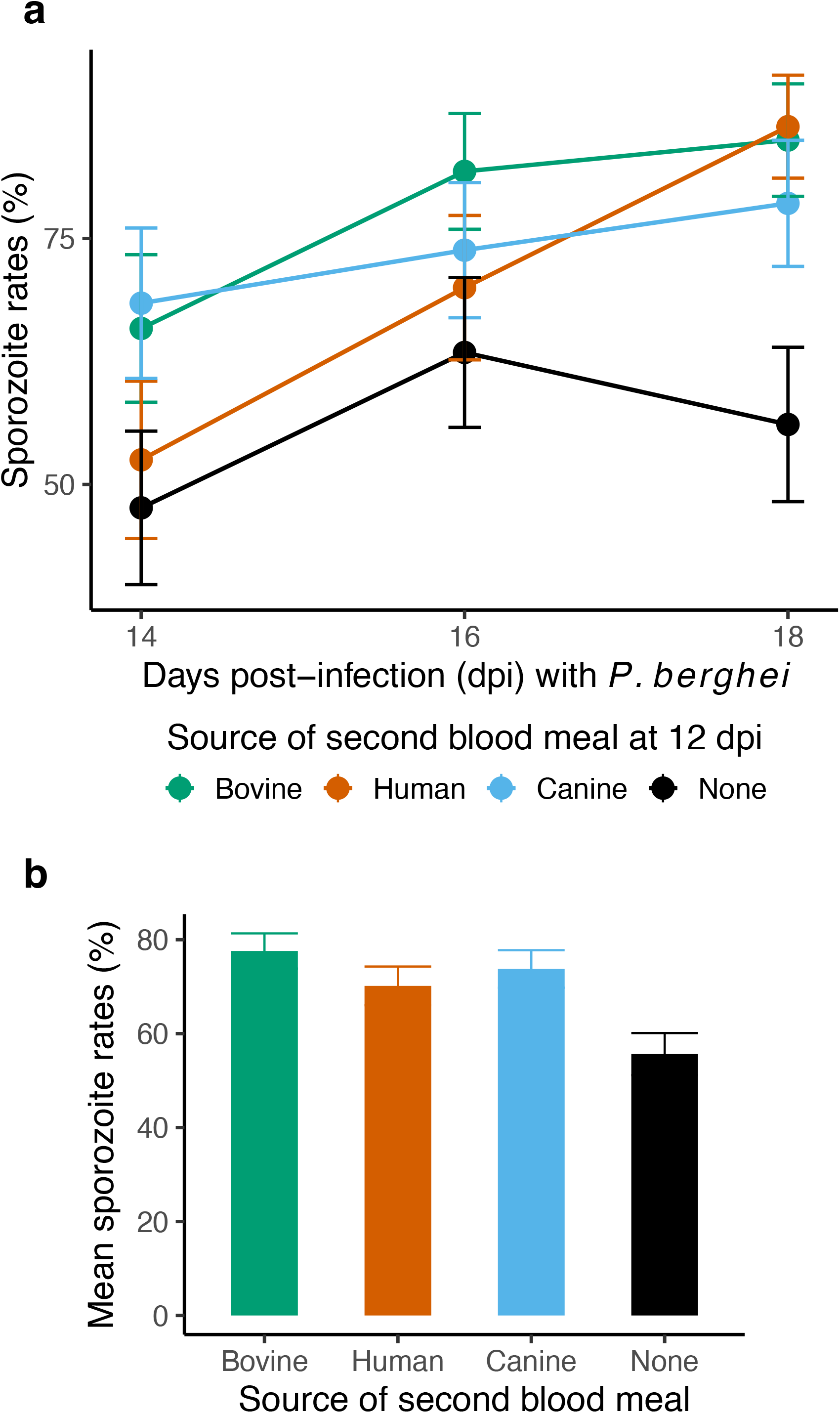
After the initial infectious blood meal, a second blood meal at 12 days post-infection from bovine (green lines/points/columns), human (red) and canine (blue) donors, (**a**) enhances migration of rodent-specific *P. berghei* sporozoites to the salivary glands, and (**b**), the mean proportion of infected mosquitoes, compared to mosquitoes that were not offered a second blood meal (‘None’, black lines/columns). Data is presented as the mean and standard errors in the proportion of sporozoite-positive mosquitoes from total sampled over two independent infections (biological replicates); in each infection, mosquitoes were offered whole blood from the same two independent donors as with *P. falciparum* infections (Figure 1). For statistical analyses, refer to Table 1.

Highest rates of sporozoite migration were noted for *P. falciparum*-infected mosquitoes offered a second human blood meal, suggesting potential for parasite host-specificity in selectively enhancing EIP (Figure 1) (Table 1). As such, it is possible that mouse blood may selectively modify the EIP of mosquitoes previously infected with the rodent-specific *P. berghei* and routinely passaged in mice [25]. Due to logistical constraints, however, we were only able to test blood from this host in one replicate (Replicate 3, Supplementary figure 3). Statistical analysis suggested a marginal increase in sporozoite migration rates over time (OR = 1.78, 95%CI = 1.00 – 3.17, p =0.049); compared to the control groups (‘None’) however, no differences in overall (mean) rates of migration were noted for human or murine blood fed groups (p>0.05, remaining output not shown). Pairwise comparisons of the model fitted estimates suggested higher prevalence of sporozoite positive mosquitoes after human blood than murine blood (estimate = 0.281, standard error = 0.09, p = 0.005).

### Blood meals from distant vertebrate hosts and survival of *Plasmodium*-infected mosquitoes

For *P. falciparum*-infected mosquitoes, survival was assessed daily from 8-21 days for all the treatments in each of the two replicates. Survival of mosquitoes in the control groups (‘None’) was indistinguishable from the groups offered human (Risk Ratio (RR) = 0.98, 95%CI = 0.57 – 1.68, p=0.949) or bovine blood (RR = 0.96, 95%CI = 0.56 – 1.65, p=0.885) from either donor (Figure 3a) (Supplementary table 4). However, both donors of canine blood were toxic to mosquitoes (Risk Ratio = 6.48, 95%CI = 4.25 – 9.85, p<0.001), with the effect particularly apparent in the first two days following the second blood meal (Figure 3a).

**Figure 3.**
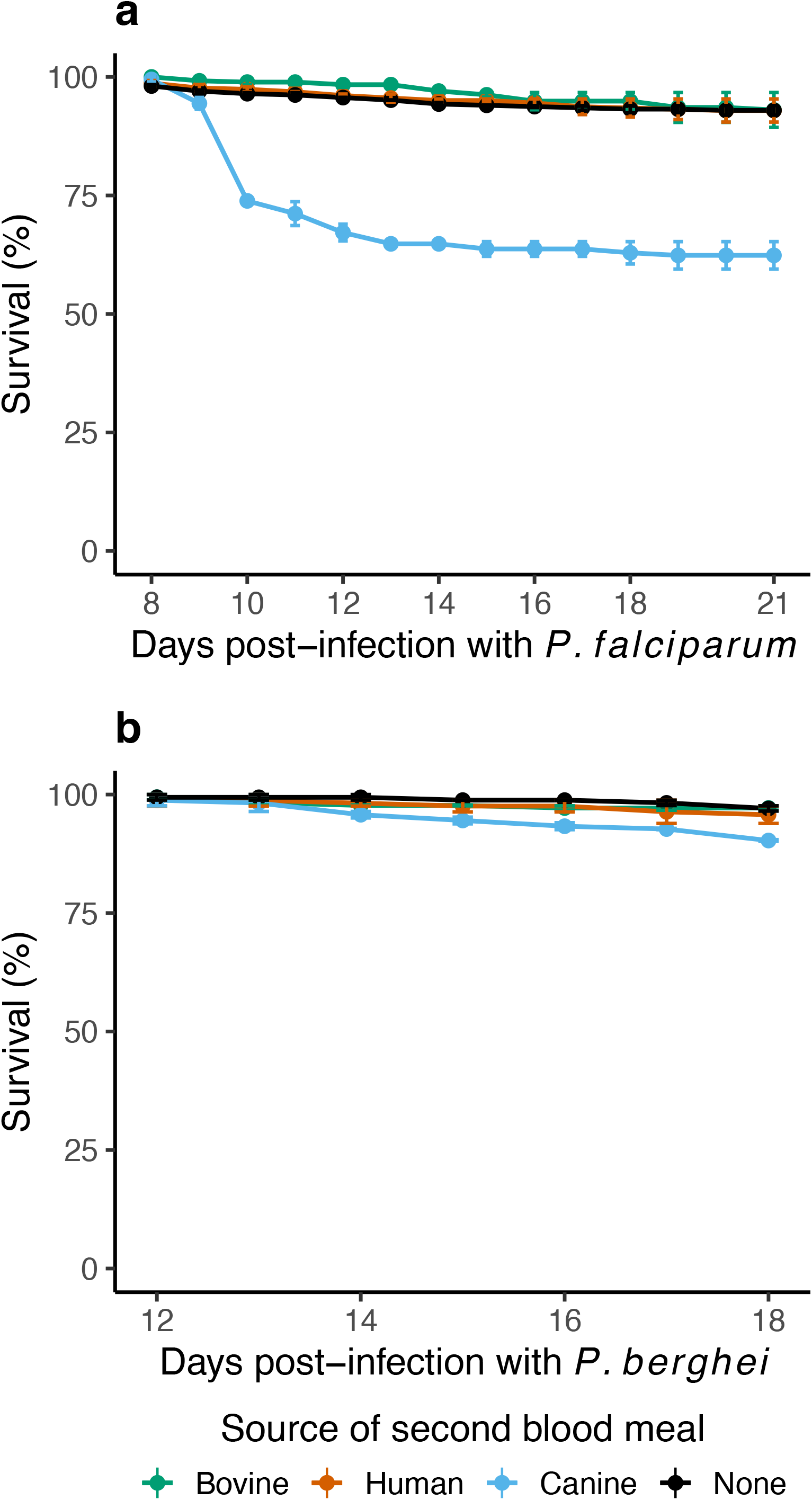
Survival rates of mosquitoes challenged with (**a**) a *P. falciparum*-spiked blood meal and (**b**) *P. berghei-*infected mice, and then offered a second blood meal bovine (green lines/points), human (red) and canine (blue) donors, compared to mosquitoes that were challenged with parasite but not offered a second blood meal (‘None’, black). Data was summarized as the mean and standard errors from two independent infections (biological replicates), where whole blood from two independent donors from all sources was offered to the mosquitoes in each infection. For statistical analyses, refer to Supplementary table 4.

For *P. berghei*-infected mosquitoes in replicates one and two, survival was assessed daily between 12-18 days post-infection; survival was not measured in replicate three. While human or bovine blood meals did not alter mosquito survival (Figure 3b) (Supplementary table 4), canine blood reduced survival of mosquitoes (RR = 3.47, 95%CI = 1.27 – 9.48, p = 0.015) (Figure 3b), although not to the same extent as the *P. falciparum*-infected groups above (Figure 3a).

### Can choice of anticoagulant confound measurements of mosquito survival?

Our results above suggested canine blood was associated with lower rates of survival for *P. falciparum*-exposed mosquitoes and to a lesser, yet clear extent for *P. berghei*-exposed mosquitoes (Figure 3) (Supplementary table 4). While it is possible that canine blood alone was toxic to the mosquitoes, another limitation to consider was the anticoagulant, with a few studies showing interaction with blood components influencing vector life-history traits, including survival [29-31]. Since we did not compare survival of *Plasmodium*-challenged groups simultaneously with naïve mosquitoes (not exposed to *Plasmodium*), we performed separate experiments to assess whether anticoagulants were confounding factors. Naïve mosquitoes were offered whole blood from various vertebrate species collected into heparin as above, or EDTA, another commonly used anti-coagulant (Figure 4). For mosquitoes housed in the same conditions as *P. falciparum*-challenged mosquitoes above, compared to non-blood fed mosquitoes, mosquitoes fed canine blood collected in EDTA were at elevated risk of mortality (RR = 3.68, 95%CI = 2.72 – 4.99, p<0.001) (Figure 4a, blue line) (Supplementary table 5). By contrast, the risk of mortality was lower for mosquitoes offered canine blood collected in heparin (RR = 0.48, 95%CI = 0.33 – 0.69, p<0.001), or bovine blood collected in EDTA (RR = 0.48, 95%CI = 0.34 – 0.68, p<0.001) or heparin (RR = 0.42, 95%CI = 0.29 – 0.61, p<0.001) suggesting that mortality associated with canine blood meals was confounded by the anticoagulant.

**Figure 4.**
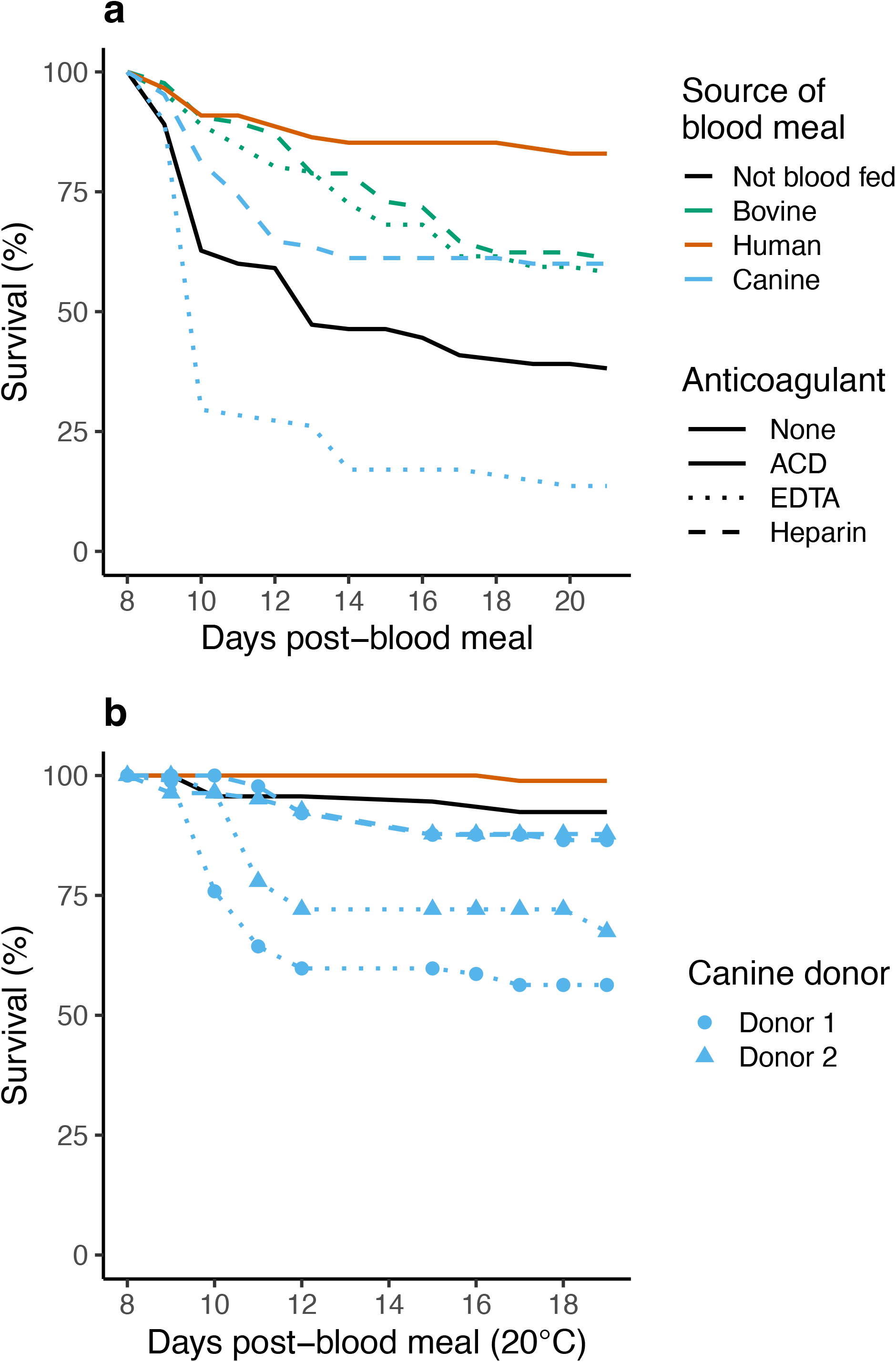
To determine whether the toxicity noted with canine blood was due to the anticoagulant, survival rates of naïve (not *Plasmodium* challenged), non-blood fed mosquitoes (‘Not blood fed’ / ‘None’, black lines) was compared to groups offered whole blood from the various sources (bovine (green)/human (red)/canine (blue)) collected in the anticoagulants citrate-dextrose (‘ACD’, solid lines, human only), heparin (dashed lines) or Ethylenediaminetetraacetic acid (‘EDTA’, dotted lines), (**a**), to replicate experimental conditions for *P. falciparum*-challenged mosquitoes (see Figure 1 and Figure 3a), and (**b**), 20°C to replicate experimental conditions for *P. berghei*-challenged mosquitoes (see Figure 2 and Figure 3b). Data is from one replicate representing each condition and thus, standard errors were not estimated. Other differences of note compared to data from *Plasmodium*-infected mosquitoes (Figures 1, 2, and 3) are, 1) survival rates were assessed after a single blood meal, and 2), in (**b**), toxicity of anticoagulants was assessed with two separate canine donors (blue filled circles and triangles). For statistical analyses, refer to Supplementary table 5.

This confounding effect was further confirmed for mosquitoes housed at 20°C (to mimic conditions for *P. berghei*), albeit with two independent canine blood donors (Figure 4b) (Supplementary table 5). Compared to non-blood fed mosquitoes, groups fed canine blood collected in EDTA were at higher risk of mortality irrespective of whether blood was collected from the first (RR = 4.68, 95%CI = 2.22 – 9.85, p<0.001) or the second donor (RR = 7.02, 95%CI = 3.40 – 14.52, p<0.001). Canine blood collected in heparin was not associated with the same risk as EDTA when blood was collected from the first (RR = 1.70, 95%CI = 0.73 – 3.98, p=0.221) or second donor (RR = 1.83, 95%CI = 0.80 – 4.18, p=0.152), or if human blood was collected in ACD (RR = 0.64, 95%CI = 0.23 – 1.81, p=0.405); note that we were unable to test the effect of bovine blood in this experiment. Taken together, our results suggest canine blood collected in either anticoagulant was toxic even to naïve mosquitoes, although compared to EDTA, the toxicity associated with heparin was significantly lower.

## DISCUSSION

Additional blood meals offered to mosquitoes reduced the EIP (time to infectiousness) and/or overall proportion of mosquitoes infected with human and rodent *Plasmodium* species. In the absence of any apparent preference for a particular blood source, human blood strongly augmented the development of the *P. falciparum*, with bovine blood showing the strongest enhancement for *P. berghei*. However, it was unclear whether the enhancement with *P. falciparum* reflected host specificity because sporozoite rates for the rodent-restricted *P. berghei* were not enhanced by mouse blood. In general, however, the additional blood meals shortened the EIP for *P. falciparum*, while also enhancing the overall proportion of *P. berghei*-infected mosquitoes. Blood from human and bovine donors was more consistent in their enhancement than canine blood, although the latter’s effect may have been confounded by its toxicity to mosquitoes; follow-up experiments with naïve mosquitoes partly recapitulated this result, suggesting that the toxicity was an artifact of anti-coagulant rather than canine blood per se. Taken together, although our results highlight the limitations of a laboratory system, they also suggest that in natural settings, where mosquitoes feed on live hosts, the potential for any non-human blood meal to accentuate the risk of human infections by enhancing development rates of any *Plasmodium* species, cannot be ignored.

Additional blood meal(s) can reduce the EIPs of several vector-borne pathogens [2-4]. A recent study with *An. coluzzi* infected with *P. falciparum* or *P. berghei* suggested the reduced EIP may be due to accelerated sporozoite replication and migration from the oocysts after the additional blood meal [32]. While this enhancement may be dependent on when (i.e., days post-infection) the additional blood meal was provided, in general, additional blood meals offered between 3-11 days post-infection all resulted in reduced EIP (reviewed by [3]). While enhancement was also observed for the rodent-specific *P. berghei*, sporozoite migration was already noted by 12 days post-infection (mean ± se = 14.5% ± 5%) when the mosquitoes were offered the second blood meal and as such, we were unable to estimate the true EIP [2]. However, a recent study with *P. berghei*-infected mosquitoes suggested a second blood meal at 4, but not 8 days after the initial infection resulted in reduced oocyst densities, but not when offered after 8 days [33]. While the study did not address downstream consequences on sporozoite prevalence or densities, it indicates timing of blood meal may have distinct effects for this parasite. It should be noted however that *P. berghei* is rather unique in its low temperature requirement (20°C) and as such, it is possible that the effect of timing may reflect the altered mosquito physiology and/or parasite development rates at this temperature [33,34]. Taken together, our results confirm the effect of an additional blood meal was independent of oocyst densities earlier in the infection (range of 3 to 94), with shorter EIPs/enhanced migration for both parasite species suggesting the potential for additional blood meals to enhance development rates of other *Plasmodium* species.

Our colony of *An. stephensi* is propagated with human blood on a routine basis. Despite this, the mosquitoes did not appear to show any preference for blood from a particular vertebrate host (as indicated by the gravid status of individuals after the second blood meals). Although *An. stephensi* is a well-known vector of human malaria, numerous field studies suggest bovids remain the preferred host [18,35]. Indeed, our mosquito line appears to have retained its intrinsic (genetic) ability to ‘digest’ bovine blood [14]. Moreover, in comparison to canine blood (see below), additional evidence of this innate ability is also provided in part by the overall lack of toxicity with bovine blood. In general, compared to other sources of blood, bovine blood is more digestible due to its ready availability. For instance, notwithstanding the socio-economic and cultural importance of cattle to humans [20], field assays of blood feeding behavior for several species of *Anopheles* find bovine blood meals are often the most common sources of blood meals and often co-incident with human blood in the same mosquito [9,10,24]. Emami et al [6] also showed that despite being maintained on human blood, bovine blood enhanced sporozoite densities in *An. gambiae s*.*s*. (Keele) and *An. arabiensis* (Ifakara) mosquitoes infected with *P. falciparum* NF54.

In contrast to bovine blood, the association between canine and human blood in mosquitoes is less evident, especially for *An. stephensi* [10]. In general, however, surveys from various geographical regions show *Anopheline* mosquitoes can feed, or even prefer canine blood in certain circumstances [11,36-38]. However, few studies have assessed whether these blood meals were coincident with human *Plasmodium* infections in mosquitoes [9,39,40]. Our results suggest canine blood may also enhance parasite development rates. While similar rates of survival were observed if blood from a bovine donor was collected in heparin or EDTA, canine blood collected in heparin, but especially EDTA showed a toxic effect on our mosquitoes. Overall toxicity was higher for *P. falciparum*-challenged mosquitoes than *P. berghei*-infected, although this have been exacerbated by housing the former group of mosquitoes at the higher temperature [41]; moreover, this temperature-dependence in toxicity was also noted with naïve mosquitoes (Figure 4).

Although toxicity was lower with heparin as the anti-coagulant, it was still higher than the other blood sources. As such, it is possible that *Plasmodium*-infection may have exacerbated toxicity; in other words, our results do not exclude the possibility of the toxic product(s) selectively targeting infected or un-infected (*Plasmodium* exposed but not infected). We speculate this is unlikely because 1) similar rates of mortality were noted for heparinized canine blood provided to naïve mosquitoes and 2), at 20°C where toxicity associated with the anti-coagulant was negligible compared to the higher temperature (Figure 4), canine blood was still consistent in its enhancement potential (Figure 2), thus supporting our overall conclusion that canine blood also enhanced parasite development rates in mosquitoes.

In addition to some of the caveats listed above, we suggest another three worth considering. First, although we did not compare survival of *Plasmodium*-challenged groups in parallel with naïve mosquitoes (not exposed to *Plasmodium*), taken together, our results underscore the importance of testing anticoagulants before initiating any empirical investigation in mosquito blood feeding behavior. Future experiments with a nontoxic anti-coagulant (e.g., Citrate-Dextrose or ACD) may reveal the true extent of its effect, although our results also indicate that the effects may not be ‘linear’ and are likely to be confounded by complex underlying interactions. For instance, while some blood sources were better at enhancing parasite development rates, it is unclear to what extent these were influenced by anticoagulant, in addition to the effect of temperature on the various measures [41-43]. It is also worth noting that these effects may be restricted to *Anopheles* as in general, anticoagulants appear to be less toxic for *Culex* and *Aedes* species [29-31]. Second, although our mosquitoes did not appear to show any preference for a particular source of blood, our measure of feeding rates qualitative (gravid/not gravid, supplementary figure 2) and as such, does not reflect the rates of digestion; indeed, the authors (JCS and AKP) noted delayed digestion of blood in midguts as well as slower rates of egg synthesis with canine, but not human or bovine blood. While this lack of ‘digestibility’ could also explain why no enhancement was seem with murine blood in *P. berghei*-infected mosquitoes (Supplementary figure 3), in general, such observations have been quantified previously with differences noted between source of blood as well as *Anopheles* species [1]. As such, it is possible the slower rates of digestion of canine blood may be responsible for the delayed migration of sporozoites to the salivary gland (Figure 1). Third, although we tested the effect of a single additional blood meal, both number and timing of the additional blood meals may alter the EIP [2,3]. A recent study with *P. falciparum* showed timing of the additional blood meal altered sporozoite densities: compared to the control groups, highest densities were noted after a blood meal at 3 days post-infection, followed by 6 and 9 days respectively. While this suggests our experimental design may have underestimated the effects of the bloodmeals on EIP, it should not affect our overall conclusions.

## CONCLUSIONS

Our results suggest how vector ecology can directly influence parasite transmission, with implications for policy. Together with the study by Emami et al [6], a major implication of our findings is that in addition to the positive effect of non-human blood meals on population density of vectors [16], they may directly impact parasite development and transmission potential. While the increasing failure of ITNs is partly attributed to the emergence of vector species with mixed host preferences [18], a recent study from Burkina Faso showed sporozoites rates in wild-caught mosquitoes were unexpectedly high for a region with low human blood indices (<20%) [12,13]; our results suggest non-human blood meals should be re-evaluated as a risk factor. Whether the reduced digestibility of canine blood reflects *An. stephensi*’s innate preference for bovine blood is unclear, however, while the traits conferring preference have been shown to be heritable, meta-analyses repeatedly suggest host preferences are driven by local adaptations to host availability [1,7,24]; our results suggest local adaptation of mosquitoes may also be dependent on the density and/or biodiversity of vertebrate fauna [44]. Finally, we posit that our observations warrant confirmation with any vector with mixed host preferences, including other species of Anopheles, as well as species of *Aedes* and *Culex* that transmit arboviruses [1,45].

## ACKNOWLEDMENTS

The authors would like to thank Courtnie Vickery for help with data collection, Anne Elliot and Jared Cooper for their help with mosquito husbandry, and the staff at UGA animal resources for mouse husbandry.

## AUTHOR CONTRIBUTIONS

**Ashutosh K. Pathak**. Conceptualization; Data curation; Formal analysis; Funding acquisition; Investigation; Methodology; Project administration; Resources; Supervision; Visualization; Roles/Writing - original draft. **Justine C. Shiau:** Data curation; Investigation; Methodology; Resources; Supervision; Visualization; Writing - review & editing. **Cury Rafael Sadock de Freitas:** Investigation; Data curation; Resources. **Dennis E. Kyle:** Funding acquisition; Resources.

## FIGURE LEGENDS

**Supplementary figure 1.**
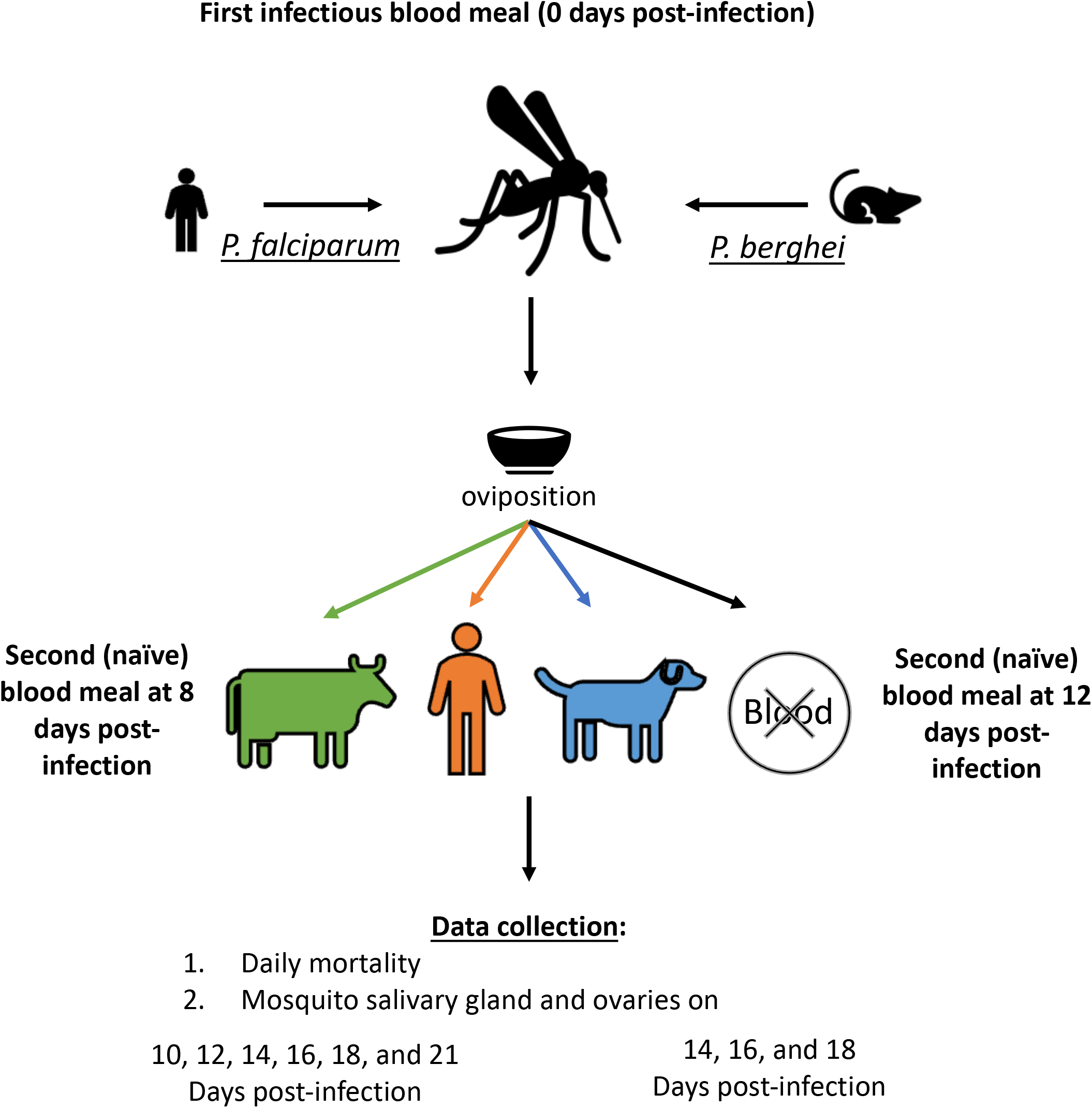
Overview of study design (for detailed description, refer to ‘Study design’ section under ‘Methods’).

**Supplementary figure 2.**
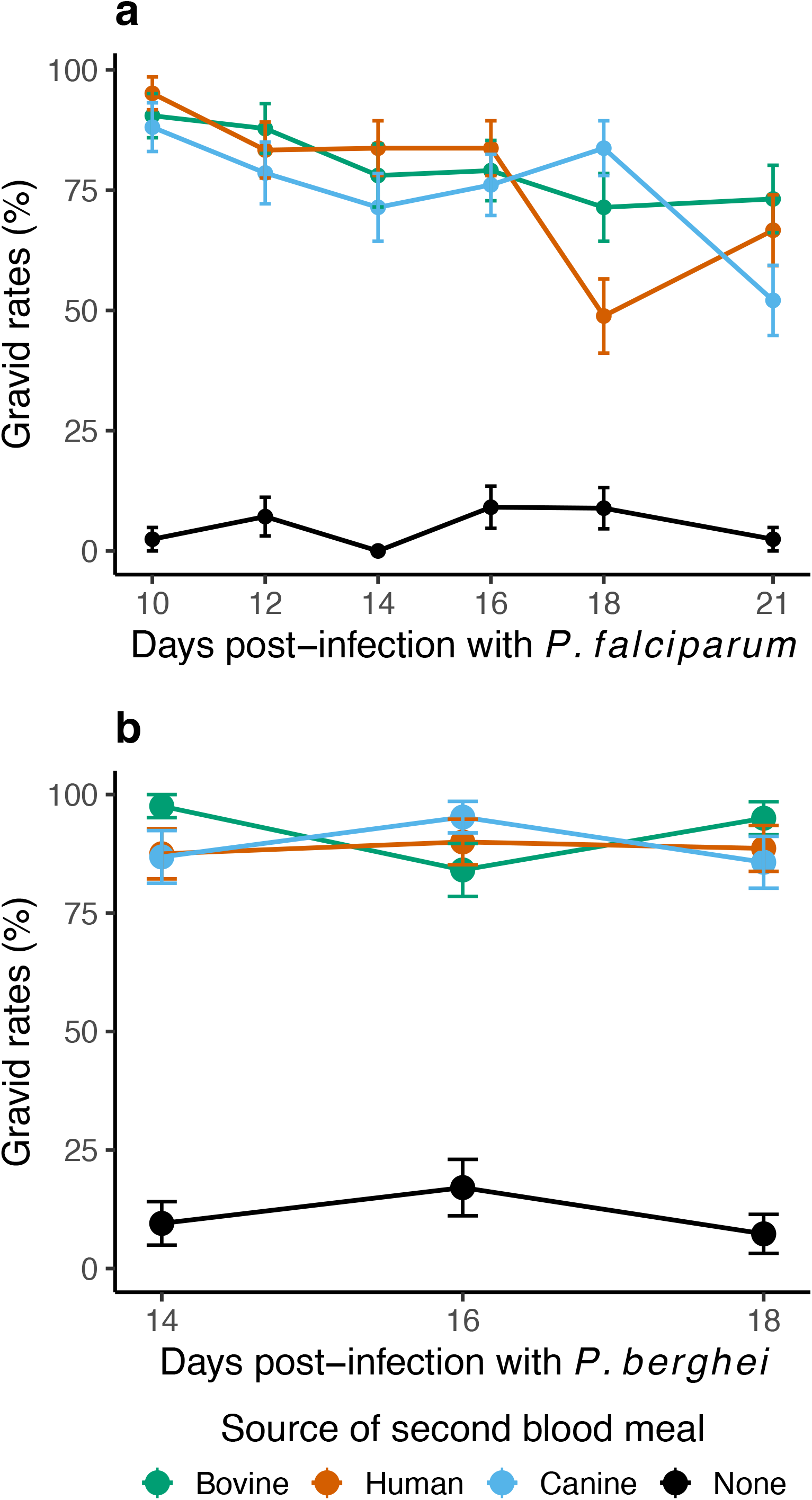
Feeding rates for the various blood meals (indicated by colors) were assessed as the proportion of mosquitoes with eggs in the ovaries (‘gravid’), following initial challenge with (**a**) *P. falciparum* and (**b**) *P. berghei*. Black lines depict the control group that were initially challenged with parasite, but not offered a second blood meal (‘None’); low rates of gravidity indicate high rates of oviposition prior to the second bloodmeal (for detailed rationale, refer to ‘Study design’ section under ‘Methods’). Note that gravid rates were measured from the same mosquitoes that were checked for sporozoite presence in the salivary glands (Figure 1 and Figure 2, also see ‘Study design’ section under ‘Methods’); as such, data represents mean and standard error from two independent replicates. See supplementary table 1 for statistical analysis and supplementary tables 2 and 3 for post hoc pairwise comparisons of the means estimated by the analyses.

**Supplementary figure 3.**
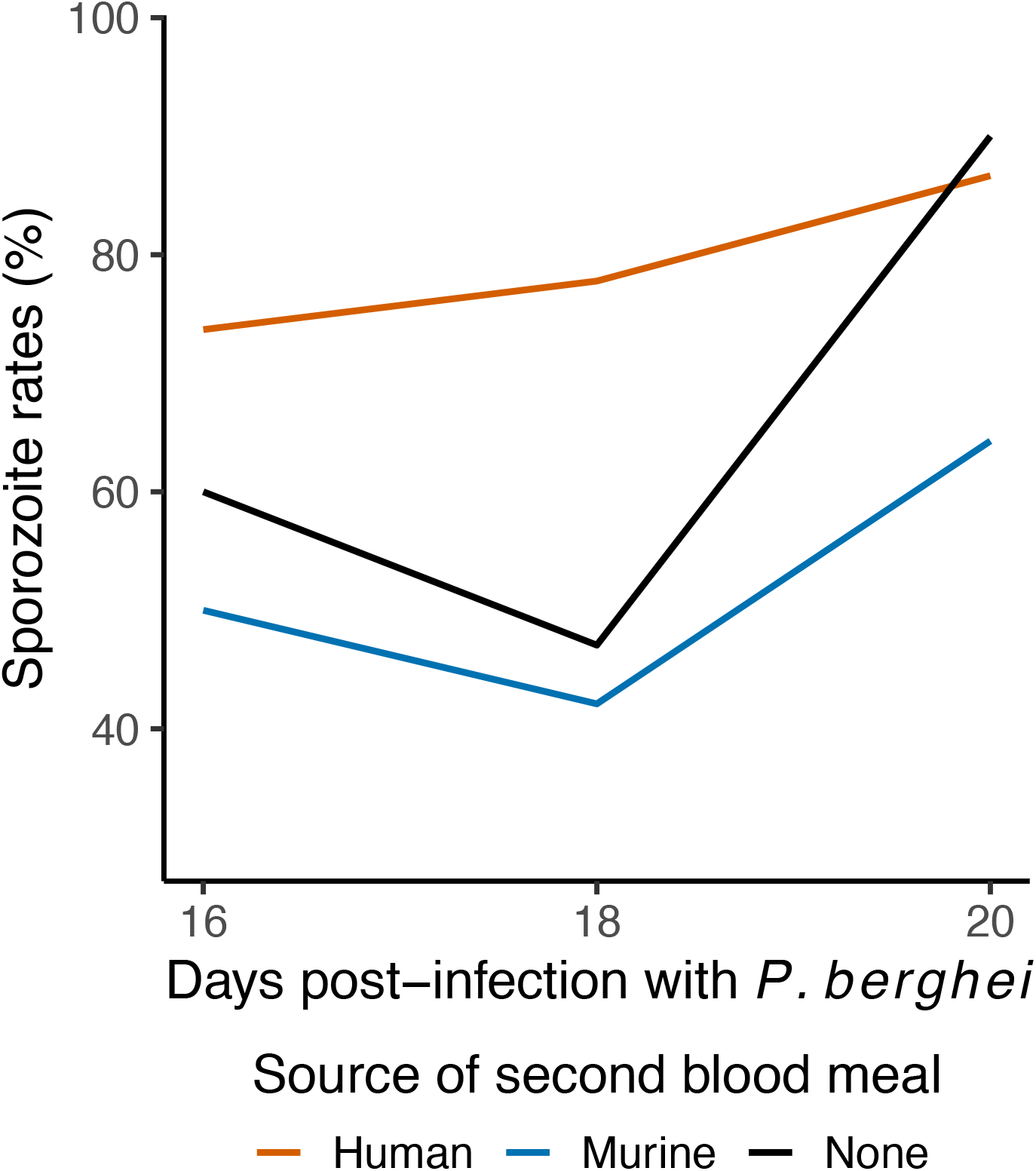
After the initial infectious blood meal, a second blood meal from human (red lines), canine (blue) and a mouse (dark blue) donor altered migration rates of rodent-specific *P. berghei* sporozoites to the salivary glands, compared to mosquitoes that were not offered a second blood meal (‘None’, black). Data is from one replicate and therefore, standard errors were not estimated.

## TABLE LEGENDS

**Supplementary table 1.** Statistical model of feeding rates for groups challenged with either parasite, to compare proportions of mosquitoes with eggs in ovaries (‘gravid’) from total sampled.

| *Predictors* | Gravid rates of *P. falciparum*<br>challenged mosquitoes |  |  | Gravid rates of *P. berghei*<br>challenged mosquitoes |  |  |
| --- | --- | --- | --- | --- | --- | --- |
|  | *Odds Ratios* | *CI* | *p* | *Odds Ratios* | *CI* | *p* |
| (Intercept) | 0.05 | 0.02 – 0.12 | **<0.001** | 0.13 | 0.07 – 0.22 | **<0.001** |
| Bovine blood<br>[vs. None] | 83.72 | 43.54 – 160.98 | **<0.001** | 91.29 | 38.75 – 215.09 | **<0.001** |
| Human blood<br>[vs. None] | 74.48 | 38.76 – 143.13 | **<0.001** | 62.01 | 28.19 – 136.42 | **<0.001** |
| Canine blood<br>[vs. None] | 62.54 | 33.00 – 118.49 | **<0.001** | 66.42 | 29.75 – 148.28 | **<0.001** |
| Days post-infection (dpi) | 1.13 | 0.64 – 1.99 | 0.670 | 0.92 | 0.53 – 1.60 | 0.758 |
| Bovine blood *<br>dpi | 0.59 | 0.31 – 1.13 | 0.110 | 0.94 | 0.40 – 2.23 | 0.890 |
| Human blood *<br>dpi | 0.44 | 0.23 – 0.84 | **0.013** | 1.14 | 0.52 – 2.49 | 0.743 |
| Canine blood *<br>dpi | 0.56 | 0.29 – 1.04 | 0.068 | 1.03 | 0.46 – 2.29 | 0.952 |
| **Random Effects** |  |  |  |  |  |  |
| $\sigma^2$ | 3.29 | | | 3.29 | | |
| T00 | 0.22 replicate |  |  | 0.01 replicate |  |  |
| Replicates | 2 |  |  | 2 |  |  |
| Observations | 1022 |  |  | 495 |  |  |

**Supplementary table 2.** Pairwise comparisons of gravid rates between the various *P. falciparum* infected groups over time (dpi = days post-infection), based on means estimated by the model in Supplementary table 1.

| **Comparison** | **Contrast - Reference** | **Dpi** | **estimate** | **SE** | **p.value** |
| --- | --- | --- | --- | --- | --- |
| 1 | None - Bovine | 10 | -0.8427245 | 0.0436962 | **<0.001** |
| 2 | None - Bovine | 12 | -0.8152954 | 0.0429059 | **<0.001** |
| 3 | None - Bovine | 14 | -0.7834499 | 0.0438422 | **<0.001** |
| 4 | None - Bovine | 16 | -0.7468820 | 0.0484963 | **<0.001** |
| 5 | None - Bovine | 18 | -0.7054399 | 0.0584838 | **<0.001** |
| 6 | None - Bovine | 21 | -0.6343493 | 0.0836175 | **<0.001** |
| 7 | None - Human | 10 | -0.8703899 | 0.0362690 | **<0.001** |
| 8 | None - Human | 12 | -0.8316284 | 0.0393372 | **<0.001** |
| 9 | None - Human | 14 | -0.7808002 | 0.0448906 | **<0.001** |
| 10 | None - Human | 16 | -0.7162754 | 0.0541697 | **<0.001** |
| 11 | None - Human | 18 | -0.6378487 | 0.0679994 | **<0.001** |
| 12 | None - Human | 21 | -0.4993026 | 0.0944482 | **<0.001** |
| 13 | None - Canine | 10 | -0.8199915 | 0.0481320 | **<0.001** |
| 14 | None - Canine | 12 | -0.7839430 | 0.0491017 | **<0.001** |
| 15 | None - Canine | 14 | -0.7416926 | 0.0516501 | **<0.001** |
| 16 | None - Canine | 16 | -0.6930624 | 0.0572830 | **<0.001** |
| 17 | None - Canine | 18 | -0.6382744 | 0.0671350 | **<0.001** |
| 18 | None - Canine | 21 | -0.5462504 | 0.0893233 | **<0.001** |
| 19 | Bovine - Human | 10 | -0.0276654 | 0.0421106 | 0.9131034 |
| 20 | Bovine - Human | 12 | -0.0163330 | 0.0401895 | 0.9773174 |
| 21 | Bovine - Human | 14 | 0.0026497 | 0.0375722 | 0.9998742 |
| 22 | Bovine - Human | 16 | 0.0306065 | 0.0388904 | 0.8603850 |
| 23 | Bovine - Human | 18 | 0.0675912 | 0.0507910 | 0.5434628 |
| 24 | Bovine - Human | 21 | 0.1350467 | 0.0869349 | 0.4059910 |
| 25 | Bovine - Canine | 10 | 0.0227330 | 0.0471544 | 0.9630661 |
| 26 | Bovine - Canine | 12 | 0.0313523 | 0.0434290 | 0.8883709 |
| 27 | Bovine - Canine | 14 | 0.0417573 | 0.0393154 | 0.7126953 |
| 28 | Bovine - Canine | 16 | 0.0538196 | 0.0394533 | 0.5222462 |
| 29 | Bovine - Canine | 18 | 0.0671655 | 0.0496484 | 0.5294257 |
| 30 | Bovine - Canine | 21 | 0.0880989 | 0.0829140 | 0.7124404 |
| 31 | Human - Canine | 10 | 0.0503984 | 0.0449650 | 0.6767417 |
| 32 | Human - Canine | 12 | 0.0476854 | 0.0431271 | 0.6861070 |
| 33 | Human - Canine | 14 | 0.0391076 | 0.0398671 | 0.7603632 |
| 34 | Human - Canine | 16 | 0.0232131 | 0.0398365 | 0.9372975 |
| 35 | Human - Canine | 18 | -0.0004257 | 0.0506656 | 0.9999998 |
| 36 | Human - Canine | 21 | -0.0469478 | 0.0861807 | 0.9479580 |

**Supplementary table 3.** Pairwise comparisons of gravid rates between the various *P. berghei* infected groups over time (dpi = days post-infection), based on means estimated by the model in Supplementary table 1.

| **Comparison** | **contrast - reference** | **Dpi** | **estimate** | **SE** | **p.value** |
| --- | --- | --- | --- | --- | --- |
| 1 | None - Bovine | 14 | -0.8091793 | 0.05847327 | **<0.001** |
| 2 | None - Bovine | 16 | -0.8080066 | 0.03742776 | **<0.001** |
| 3 | None - Bovine | 18 | -0.8039053 | 0.06028329 | **<0.001** |
| 4 | None - Human | 14 | -0.7578363 | 0.06618743 | **<0.001** |
| 5 | None - Human | 16 | -0.7744462 | 0.04022415 | **<0.001** |
| 6 | None - Human | 18 | -0.7899469 | 0.0605715 | **<0.001** |
| 7 | None - Canine | 14 | -0.7771955 | 0.06397787 | **<0.001** |
| 8 | None - Canine | 16 | -0.7812983 | 0.03988145 | **<0.001** |
| 9 | None - Canine | 18 | -0.7840726 | 0.06215741 | **<0.001** |
| 10 | Bovine - Human | 14 | 0.05134306 | 0.05838495 | 0.81556261 |
| 11 | Bovine - Human | 16 | 0.03356039 | 0.03742676 | 0.80656636 |
| 12 | Bovine - Human | 18 | 0.01395843 | 0.06030173 | 0.99562756 |
| 13 | Bovine - Canine | 14 | 0.0319838 | 0.05582956 | 0.94014262 |
| 14 | Bovine - Canine | 16 | 0.02670826 | 0.03704534 | 0.88873322 |
| 15 | Bovine - Canine | 18 | 0.01983275 | 0.06189679 | 0.98861436 |
| 16 | Human - Canine | 14 | -0.0193593 | 0.06389918 | 0.99033589 |
| 17 | Human - Canine | 16 | -0.0068521 | 0.03987993 | 0.99819555 |
| 18 | Human - Canine | 18 | 0.00587432 | 0.06216987 | 0.99969756 |

**Supplementary table 4.** Statistical modeling of survival rates of *Plasmodium-*infected mosquitoes.

| Supplementary table 4 |  |  |  |  |  |  |
| --- | --- | --- | --- | --- | --- | --- |
| *Predictors* | *P. falciparum* mortality risk |  |  | *P. berghei* mortality risk |  |  |
|  | *Risk ratio* | *95%CI* | *p* | *Risk ratio* | *95%CI* | *p* |
| Bovine blood [vs. None] | 0.96 | 0.56 – 1.65 | 0.885 | 0.98 | 0.28 – 3.38 | 0.973 |
| Human blood [vs. None] | 0.98 | 0.57 – 1.68 | 0.949 | 1.46 | 0.46 – 4.61 | 0.516 |
| Canine blood [vs. None] | 6.48 | 4.25 – 9.85 | **<0.001** | 3.47 | 1.27 – 9.49 | **0.015** |
| **Random Effects** |  |  |  |  |  |  |
| T<sub>00</sub> | 0.03<sub>replicate</sub> |  |  | 0.00<sub>replicate</sub> |  |  |
| Replicates | 2 |  |  | 2 |  |  |
| Observations | 1526 |  |  | 675 |  |  |

**Supplementary table 5.**
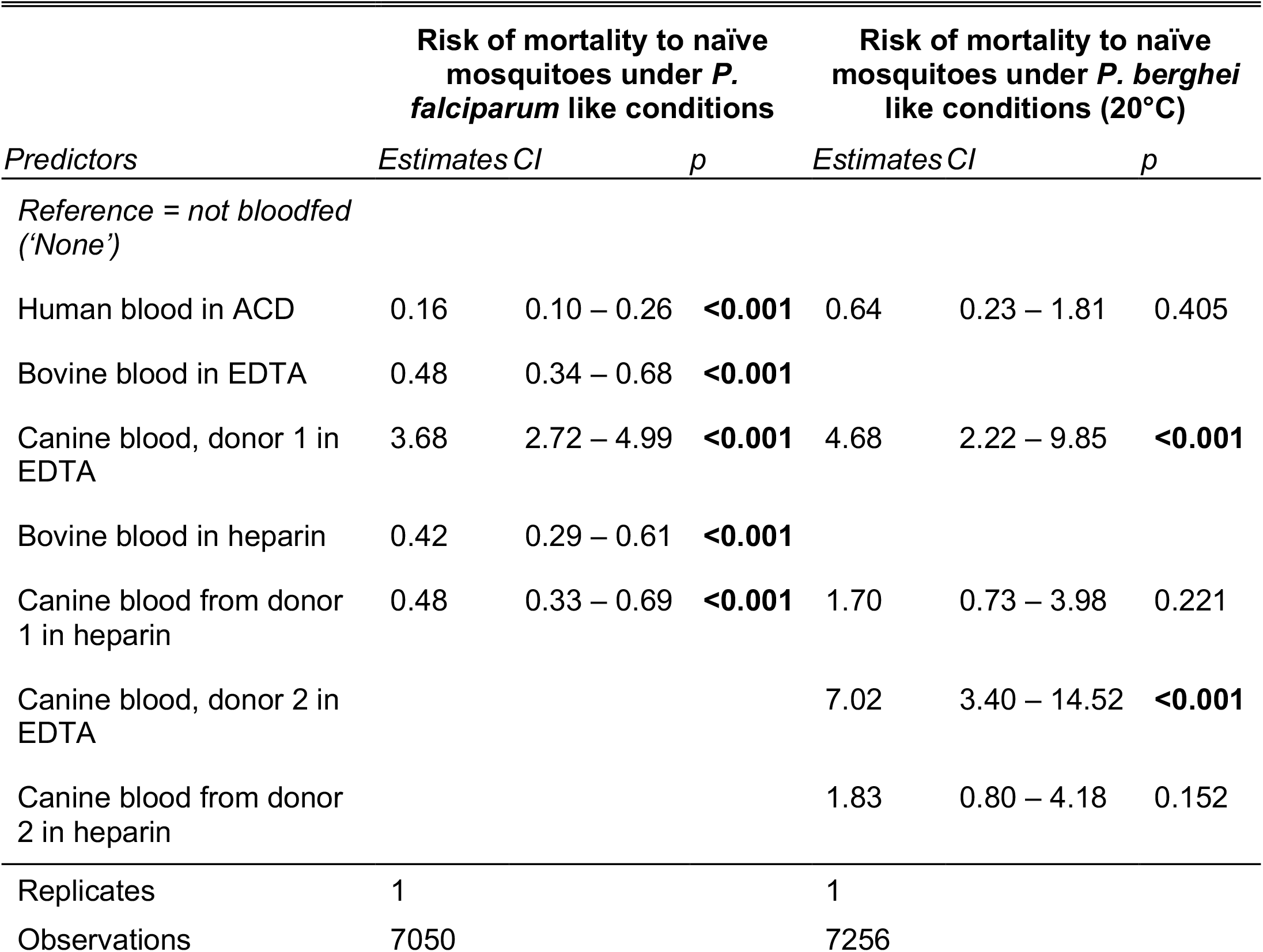
Statistical modeling of survival rates of *naïve* mosquitoes (never exposed to Plasmodium) after being offered whole blood from the various sources after collection in different anticoagulants (ACD = anticoagulant citrate-dextrose, EDTA = Ethylenediaminetetraacetic acid).

